# The 16S microbiota of Budu, the Malaysian fermented anchovy sauce

**DOI:** 10.1101/2020.03.10.986513

**Authors:** Muhammad Zarul Hanifah Md Zoqratt, Han Ming Gan

## Abstract

Budu is a Malaysian fermented anchovy sauce produced by immersing small fishes into a brine solution for 6 to 18 months. Fermentation of the anchovy sauce is contributed partly by microbial enzymes, but little is known about the microbial community in Budu. Therefore, a better understanding of the Budu microbiome is necessary to better control the quality, consistency and safety of the product. In this study, we collected 60 samples from twenty bottles of Budu produced by seven different manufacturers. We analyzed their microbiota based on V3-V4 16S rRNA amplicon sequencing at the time of opening the bottle as well as 3- and 7-months post-opening. *Tetragenococcus* was the dominant genus in many samples, reaching a maximum proportion of 98.62%, but was found in low abundance, or absent, in other samples. When Budu samples were not dominated by a dominant taxa, we observed a wider genera diversity such as *Staphylococcus, Acinetobacter, Halanaerobium* and *Bacillus*. While the taxonomic composition was relatively stable across sampling periods, samples from two brands showed a sudden increase in relative abundance of the genus *Chromobacterium* in the 7^th^ month. Based on prediction of metagenome functions, non-*Tetragenococcus*-dominated samples were predicted to have enriched functional pathways related to amino acid metabolism and purine metabolism compared to *Tetragenococcus-*dominated microbiome; these two pathways are fundamental fermented quality and health attributes of fish sauce. Within the non-*Tetragenococcus*-dominated group, contributions towards amino acid metabolism and purine metabolism were biased towards the dominant taxa when species evenness is low, while in samples with higher species evenness, the contributions towards the two pathways were predicted to be evenly distributed between taxa.

## 1 Introduction

Budu is a Malaysian fermented anchovy sauce which is prepared by immersing anchovies in brine solution (salt concentration 21.5 – 25.7 %) containing natural flavor-enhancing ingredients and left to ferment in earthen containers for 6 to 12 months (Rosma et al., 2009). Compared to other Southeast Asian fish sauces such as Nuoc Mam from Vietnam and Nampla from Thailand which are transparent, Budu is turbid and heterogenous (Lopetcharat et al., 2001)□. Fish sauce such as Budu is as food condiment because of its pleasant umami flavor. It is also nutritious because it is rich in antioxidants, vitamins and fibrin-clotting inhibitors (Shivanne Gowda et al., 2016)□. However, certain metabolites in Budu might present as a potential health risk. For instance, Budu has a high amount of purine (Li et al., 2019)□ and histamine (Rosma et al., 2009), metabolites that are associated with gout (Paul and James, 2017)□ and scombroid poisoning (Tortorella et al., 2014)□, respectively.

Fish sauce microbes are responsible towards health and gastronomic properties of the fish sauce by altering the metabolite content of the fish sauce. For instance, *Tetragenococcus muriaticus* was known to produce histamine, while *Staphylococcus, Bacillus* and *Lactobacillus* were known to express histamine oxidase which can degrade histamine (Shivanne Gowda et al., 2016)□. Therefore, a better understanding of the Budu microbial community can elevate the Budu organoleptic and health value by harnessing its microbiome. So far, all microbiological studies of Budu to date were done based on culture-based methods. For example, Yuen et al. (2009) discovered bacterial succession takes place in Budu fermentation from *Micrococcus-*to *Staphylococcus arlettae-*dominated community, and uncovered the presence of yeast species such as *Candida glabrata* and *Saccharomyces cerevisiae*.

Subsequent microbiological studies on Budu revealed a few other cultivable bacteria such as *Bacillus amyloliquefaciens* FS-05 and *Lactobacillus planatarum* which were able to produce glutamic acid which is associated with the the umami taste (Zaman et al., 2010; Zareian et al., 2012). A recent study on Malaysian fermented foods also displayed the potential of *Bacillus* sp. in producing biosurfactants that inhibit pathogenic bacterial growth (Mohd Isa et al., 2020). While culture-dependent methods are useful in gaining the first insights of role of microbes in Budu fermentation, they are biased towards microbes that can grow optimally under culture conditions. To date, there has yet any culture-independent methods to provide us an overview of the Budu microbial diversity.

In this study, we surveyed the microbial community of 60 samples from twenty bottles of Budu purchased of multiple brands. We investigated their microbial community structure at different sampling periods. We then find association between the microbial composition and diversity found in Budu with its predicted metabolic pathways. Lastly, we explored the use of microbial interaction networks modeled on 16S abundance information.

## 2 MATERIALS AND METHODS

### 2.1 Sample collection

Twenty bottles of Budu from different brands were purchased from shops in the state of Kelantan, Malaysia, at five different time points (July 2016, October 2016, November 2016, March 2017 and April 2017). These twenty bottles were sampled at three different periods (24th April 2017, 24th July 2017 and 20th November 2017), amounting a total of 60 samples. The samples were stored at room temperature to emulate typical retail storage condition. The sample metadata is shown in Table 1. Samples were named according to the following format:<brand><bottle number>_<months since last opened>.

**Table 1:** Sample metadata of Budu from seven different brands

### 2.2 DNA extraction and sequencing

200 µl of the sample was poured into 1.5 ml Eppendorf tubes and centrifuged at 14 800 rpm for 15 minutes to remove the supernatant. The samples were then resuspended in 400 µl of 20 mM Tris-EDTA buffer and added with 100 μm silica beads (OPS Diagnostics LLC, Lebanon, NJ, USA). The samples were later subjected to bead beating using Vortex Genie 2 Mo Bio (Carlsbad, CA, USA) at 3200 rpm for 1 hour. They were later added with 20 ul of 0.5% SDS and 20 ul of 50 mg/ml Proteinase K and incubated at 55°C for 1 hour. The samples were added with 100 ul of saturated potassium chloride solution and were later incubated on ice for 5 minutes. Total DNA was extracted using chloroform-isopropanol precipitation and purified with Agencourt AMPure XP beads (Beckman Coulter, Inc., Indianapolis, IN, USA).

The V3-V4 region of the 16S rRNA gene was amplified using the forward primer 5’-TCGTCGGCAGCGTCAGATGTGTATAAGAGACAGCCTACGGGNGGCWGCAG-3’ and reverse primer 5’ GTCTCGTGGGCTCGGAGATGTGTATAAGAGACAGGACTACHVGGGTATCTAATCC-3’ containing partial Illumina Nextera adapter sequence (Klindworth et al., 2013) following the Illumina 16S Metagenomic Sequencing Library Preparation protocol. To enable multiplexing, 16S amplicons were barcoded using different pairs of index barcodes to prepare DNA libraries. DNA libraries were normalised, pooled, denatured and sequenced on the Illumina MiSeq (Illumina, San Diego, CA, USA) in Monash University Malaysia Genomics Facility, using either 2 × 250 bp or 2 × 300 bp run configuration.

### 2.3 Sequence data analysis, phylogenetic tree construction, taxonomic assignment and generation of feature table

The forward and reverse PCR primer sequences were trimmed off using Cutadapt version 1.16 with default parameters (Martin, 2011). Paired-end sequences were merged using fastq_mergepairs, and quality-filtered using fastq_filter (-fastq_maxee: 1.0, -fastq_minlen: 300) in USearch v10.0.240_i86linux32 (Edgar and Flyvbjerg, 2015). High-quality sequences were then denoised to create amplicon sequence variants (ASVs); the ASV sequences can be obtained from Supplementary File 1. The ASV sequences that were generated were used as reference sequences to create a raw feature table, using UNoise3 in USearch (Edgar, 2016). The raw feature table was further processed by filtering out chloroplast and mitochondrial ASVs (see below for a description on taxonomic assignment) and by rarefying to 6,000 reads per sample for downstream analyses (Supplementary File 3).

Multiple sequence alignment of the ASV sequences was conducted using MAFFT while masking unconserved and highly gapped sites (Katoh and Standley, 2013). A phylogenetic tree was then constructed from aligned ASV sequences and was rooted at midpoint using FastTree version 2.2.10 (Bolyen et al., 2019; Price et al., 2010). 16S V3-V4 Naive-Bayes classifier was trained on V3-V4-trimmed 16S sequences of the SILVA 132 release, using q2-classifier plugin in QIIME 2 (Bokulich et al., 2018; Pedregosa et al., 2011). The SILVA reference sequences were trimmed using the same primer sequences and parameters for the raw sequencing data. The ASV sequences were then taxonomically assigned using the trained classifier; the taxonomic assignments of all ASV sequences can be found in Supplementary File 2.

### 2.4 Calculation of alpha and beta diversity indices and comparison of genera distribution between sampling batches

Species richness indices (observed ASVs and Faith PD) as well as species evenness indices (Simpson and Shannon) were calculated using QIIME2 (Bolyen et al., 2019; Hunter, 2007). All calculations of Alpha diversity indices were done at ten iterations per sequencing depth. Wilcoxon paired tests were done to compare species richness between sampling batches, as implemented using DABEST-python library (Ho et al., 2019). Since there is only one bottle for brand F and G, samples from these brands were not included in statistical comparisons of alpha diversity analyses.

Beta diversity was measured based on Bray-Curtis and weighted UniFrac to calculate distances between microbiota of the samples (Lozupone et al., 2011). ASV-and genus-level relative differential abundance between sampling batches were done using QIIME2 plugin ANCOM (Mandal et al., 2015).

### 2.5 Functional and pathway prediction

Using the normalised feature table as input data, function and pathway prediction pipelines were conducted using PiCrust2 (Douglas et al., 2020)□. Briefly, ASVs were phylogenetically placed into a reference phylogenetic tree using EPA-NG and gappa (Barbera et al., 2019; Czech et al., 2020)□. Hidden state prediction to predict 16S copy number was applied using castor R package (Louca and Doebeli, 2018)□ to normalise the feature table based on 16S copy number information. This is followed by prediction of Nearest Sequenced Taxon Index (NSTI) score per ASV. ASVs with NSTI score which is higher than 2.0 were assumed to not have a representative genome in the reference phylogenetic tree, thus were filtered out from subsequent analyses. Afterwards, prediction of gene family abundance was done against the enzyme commission, EC database, followed by prediction of pathway abundance against METACYC database (Ye and Doak, 2009)□. Scores of these predictions were normalised by sequencing depth per sample (6000 reads). Predicted functional table and predicted pathway table can be obtained from Supplementary Files 4 and 5.

Predicted pathway table was subjected to Bray Curtis distance normalization, followed by multidimensional scaling (MDS) ordination (ratio transformation) using Ecopy (https://github.com/Auerilas/ecopy). Samples with high relative abundance of *Tetragenococcus* formed a distinct cluster which separated from non-*Tetragenococcus* dominated samples. To compare potential pathways that distinguishes the two groups, post-hoc t-test was applied, adjusted by false discovery rate Benjamini Hochberg; a predicted pathway with corrected p-value of below 1<10^−5^ and Cohen d effect size of at least 3 is to be considered as significantly different between *Tetragenococcus*-dominated group and non-*Tetragenococcus*-dominated group (Terpilowski, 2019; Vallat, 2018)□.

### 2.6 Microbial association network

SPIEC-EASI 0.1.4 was used to predict the microbial association network based on co-abundance (Kurtz et al., 2015). Since SPIEC-EASI applies its own normalization method, the mitochondria and chloroplast ASVs-filtered unrarefied feature table was used as the input table. ASVs were also filtered by prevalence at the minimum occurrence of 30 samples.

Meinshausen-Bühlmann neighborhood selection model was used to model the microbial interaction network at 10,000 replications (Meinshausen and Bühlmann, 2006), and at a variability threshold of 0.05% using StARS (Liu et al., 2010). R igraph package was used for the network visualization and extracting network properties (Csardi and Nepusz, 2006). To find for important nodes in the graph, degree centrality and betweenness centrality were two node centrality measures computed from the predicted network. Degree centrality weighs a node’s importance by counting the number of edges linked to the node, while betweenness centrality evaluates the geodesic distances from all node pairs that are passing through the particular node. High degree centrality suggests a role as a keystone species, while high betweenness centrality predicts its importance in maintaining the structure of the interaction network.

## 3 RESULTS

### 3.1 The Budu microbiome is inconsistent within the same brand even at high-level taxonomic composition

Firmicutes, Proteobacteria, Halanaerobiaeota, Actinobacteria, and Bacteroidetes were the top five most abundant phyla across all Budu samples (Figure 1A) with an average relative abundance of 60.26%, 22.86%, 10.60%, 3.91%, and 1.91% respectively, while other lesser phyla contributed less than 0.5%. The cumulative relative abundance of the top five most abundant phyla in each sample ranges from 95.75% to 100.00%. The relative abundance of dominant phyla were not statistically different between sampling periods (Wilcoxon paired test p > 0.4), thus the phyla composition was fairly stable across time.

**Figure 1:**
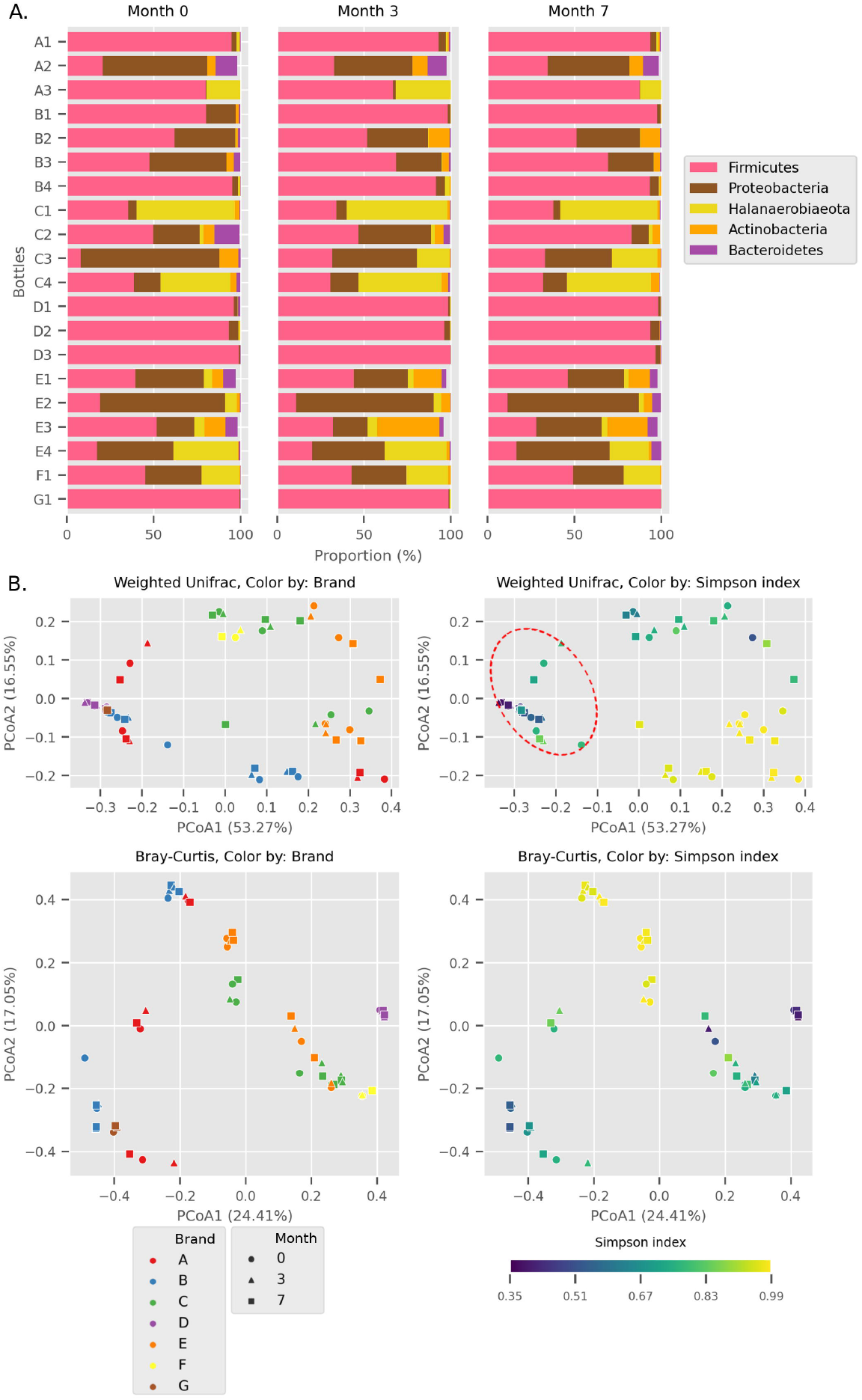
A. Relative abundance of the top five most abundant phyla in each bottle across sampling time. B. Principal Coordinates Analysis (PCoA) plots, based on Weighted UniFrac and Bray Curtis distances. Point shapes were assigned by sample batch. Point colour were by either brand, or Simpson alpha diversity index. Circular perimeter marks a cluster that was dominated by Firmicutes (> 60% relative abundance).

We observed different trends of phyla composition between and within different brands and manufacturing batches. For example, all samples from Brand D were consistently dominated by Firmicutes, making up an average relative abundance of 93% in each sample. On the contrary, some samples displayed uneven phyla distribution within brand. For example, Halanaerobiaeota made up a substantial portion from 18.8% to 48.4% in C1, C3 (except sample C3_0) and C4 samples but was markedly smaller in C2 samples (average = 2.2%). In another instance, a substantial proportion of Proteobacteria and Halanaerobiaeota was found in A2 and A3 samples respectively, which observed in very low proportion in A1 samples. The distinct phyla composition of Budu microbiome was recapitulated in the Principal Coordinates Analysis (PCoA) plot based on Weighted UniFrac distances (Figure 1B). For example, there are three clusters of brand A; each cluster consists of three points correspond to three sampling batches per bottle. This suggests that the microbiome of brand A samples from the same bottle were closely similar and that microbiome of samples of brand A of different bottles were clearly different. This sub-clustering trend was also observed in brand B, providing further evidence that the Budu microbiome observe of bottles of the same brand is inconsistent in some brands.

In weighted-UniFrac PCoA, samples that were dominated by Firmicutes cluster very closely to one side (Figure 1B; red dashed outline). This cluster included samples from multiple brands such as brand D and G. However, in the Bray Curtis distances PCoA, which weighs in only abundance information but not phylogenetic relationship between ASVs, this clustering is not observed. Brand D formed an isolated cluster away from other Firmicutes-dominated samples in Bray Curtis PCoA. This difference between weighted-UniFrac and Bray Curtis clustering indicates that brand D samples contained phylogenetically close but distinct ASVs.

Fourteen genera were found in at least half of the samples (Figure 2). Based on the enormous relative abundance (average = 38.77%, max. = 98.62%) and prevalence (96.62%), *Tetragenococcus* represents the most dominant genus of the Budu microbiome. Although it is prevalent and dominant in some samples, it was also present in low abundances in certain samples like samples from bottle A2 and samples from brand E. *Halanaerobacterium* is another genus that made up substantial proportion of some Budu microbiome (max. = 58.15%), despite its lesser prevalence than the *Staphylococcus* and *Acinetobacter. Tetragenococcus* and *Halanaerobium* are the only two genera that can reach an abundance of over 50% of reads per sample – this was observed in 21 samples (brands A, B, D and G) and 3 samples (brand C) respectively.

**Figure 2:**
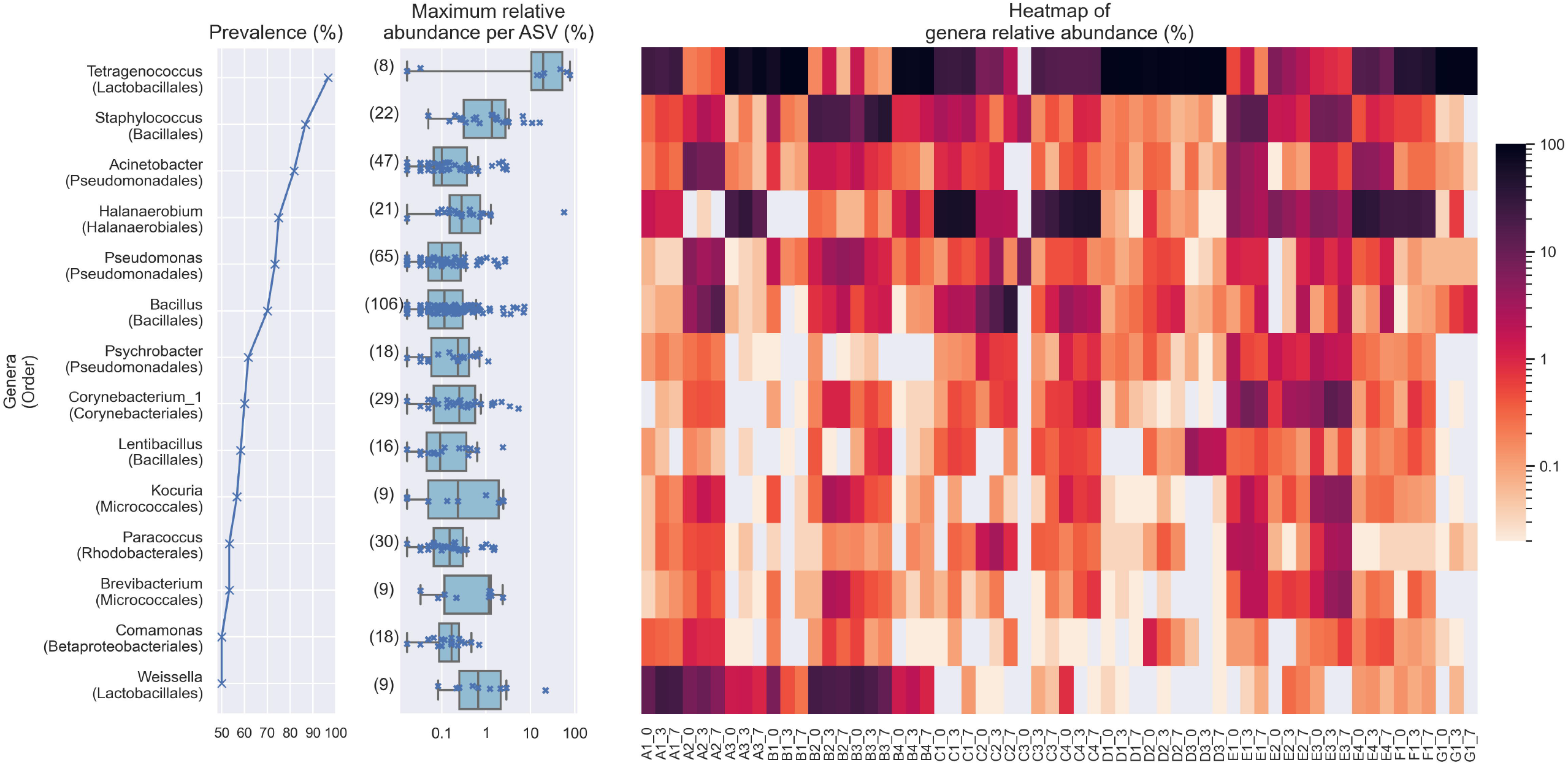
Plots indicating the prevalence of prevalent genera (at least 50% prevalence, > 0.01% abundance per sample), it’s maximum abundance for each ASV assigned to the corresponding genus and and it’s relative abundance. The bracketed numbers beside the maximum abundance per ASV boxplots indicate the number of ASVs assigned to the genus.

Despite the lower overall prevalence of *Weissella* (50%), it made over 20% of the genera composition in four samples -this degree of abundance was not observed in several genera of higher prevalence such as *Acinetobacter, Pseudomonas, Psychrobacter, Corynebacterium_1, Lentibacillus, Kocuria, Brevibacterium*, and *Paracoccus*. Meanwhile, there are other prevalent taxa which are present at lower abundance. For example, the highest abundance of *Comamonas* was at most 1.22 % of reads per sample, despite being present in half of the samples. This shows that increasing prevalence of genus in the samples is not predictive of its relative abundance. None of the core genera are 100% prevalent.

Within *Tetragenococcus*, we detected two different species which are *Tetragenococcus muriaticus* (ASV1) and *Tetragenococcus halophilus* subsp. halophilus (ASV3, ASV4 and ASV7) (Supplementary Figure 1). *T. muriaticus* composition was consistently high in brand D, while *T. halophilus* subsp. halophilus was especially abundant in some bottles of brand A, B and G. We also observed the co-presence of both *Tetragenococcus* species in some samples such as C1 samples and samples from brand F. The consistent high abundance of ASV1 in brand D is likely to explain the sub-cluster formed by brand D samples occurring in the Bray Curtis PCoA. *Halanaerobium, Staphylococcus* and *Weissella* are the only genera with ASVs above 10% relative abundance.

Bacillus was the most commonly isolated bacterial genus from Budu (Mohd Isa et al., 2020; Zaman et al., 2010)□. The abnormally enormous number of *Bacillus* ASVs (Figure 2) is possibly explained by the historically poor taxonomic demarcation of the *Bacillus* taxonomy (Patel and Gupta, 2020)□. Only recently, recent efforts proposed the revision the *Bacillus* taxonomy (Patel and Gupta, 2020)□. By maximum likelihood phylogenetic tree of partial V3-V4 16S sequences, *Bacillus* ASVs from Budu were represented in different clades within the *Bacillus* genus (Supplementary Figure 2). ASV18 was the most abundant and prevalent *Bacillus* ASV, belonging to the Cereus clade (max. = 7.05%). However, the bootstrap support values of the tree were low, ranging from 0.34 to 0.75. This indicates the limited phylogenetic resolution of partial 16S sequence that was also shown previously (Patel and Gupta, 2020)□. Some nodes were positioned into a different clade than expected, reflecting the limitations of 16S sequence as a marker to *Bacillus* evolutionary history, in a few cases. The resolution of *Bacillus* taxonomy and functional diversity in Budu can be further confirmed in the future using genomes sequences of more *Bacillus* isolates or metagenome-assembled genomes from Budu.

### 3.2 Comparison of Budu across time reveals minimal differences in alpha diversity, followed by detectable shifts in relative abundance of a few genera

Some Budu samples within a brand had greatly different species richness (Figure 3A and B), which was apparent in brands A and E. This is congruent with the inconsistent phyla composition within a brand (Figure 1). There were also apparent changes in alpha diversity across sampling batches. After 3 months, increase in observed ASVs and Faith PD were generally observed in most bottles of brand A, C and E. The alpha diversity indices decreased while a few steadily increased in some of the bottles between month 3 and month 7. There are also inconsistent shifts in species richness in brand B samples. The rarefaction plots of observed features and Faith PD indices are shown in Supplementary Figure 3. Some rarefaction curves did not completely plateue, such as those from brand D and E did not completely plateue, indicating that certain samples may contain rare taxa which were not sampled at sequencing depth of 6000 reads.

**Figure 3:**
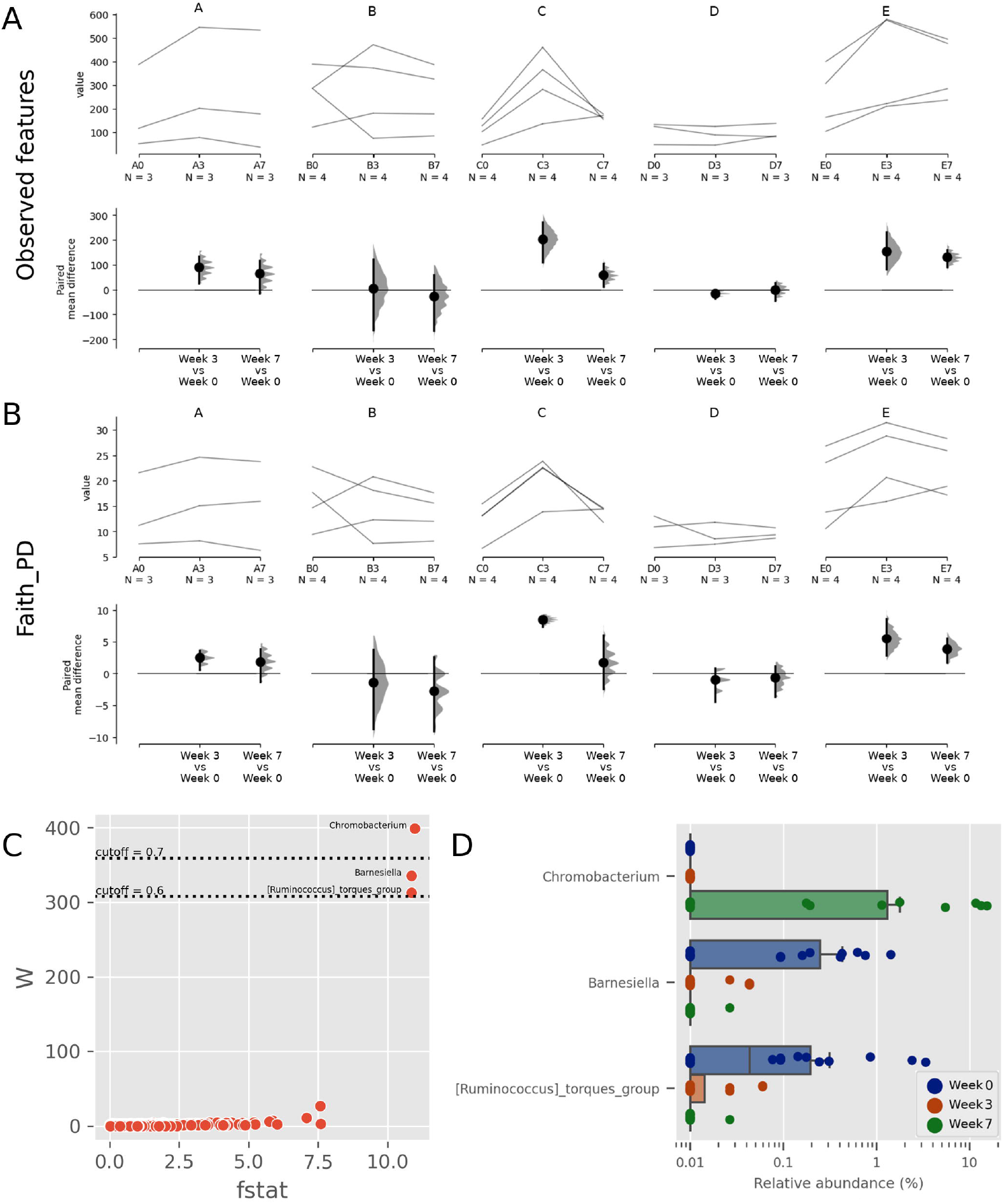
Trajectory of Observed ASVs (A) and Faith Phylogenetic Diversity (B) across sampling periods, and the comparison of alpha diversity measures at 3^rd^ and 7^th^ month sampling against control range (0^th^ Month), C) Dot plot of W statistic computed by ANCOM against F-stat of each genera, D) Relative abundance of *Chromobacterium, Barnesiella* and *[Ruminococcus] torques* group across sampling periods

When comparing the genera distribution between sampling batches, *Chromobacterium, Barnesiella* and [*Ruminococcus] torques* group displayed significantly differential abundance (W statistic > 0.6x of total features) and F statistic (> 10), as seen in Figure 3C. The relative abundance of *Chromobacterium* was higher at later sampling, reaching above 10% in brand E samples, detected in lesser abundance in brand D samples, while being virtually absent in other samples (Figure 3D, Supplementary Figure 4). The relative abundance of *Barnesiella* and *Ruminococcus torques* were much lower and did not correlate with any common metadata. Relatively minute abundances were seen in later sampling in only a few samples, suggesting that they were native members of the Budu microbiome, but at a lower relative abundance. Decreased relative abundance of these genera might be due to increase in relative abundance of other taxa. Comparison of taxonomic distribution at ASV level between sampling batches resulted in no significantly differential ASV distribution.

### 3.3 Comparison between *Tetragenococcus*-dominated and non-*Tetragenococcus*-dominated microbiome in terms of predicted pathways reveal differential enriched predicted pathways, primarily related to amino acid and purine biosynthesis

Based on predicted pathways information, samples dominated by *Tetragenococcus* form a distinct cluster away from non-*Tetragenococcus*-dominated samples (PERMANOVA p-value 0.001, Figure 4A). By looking at pathways that were significantly different between the two groups, it was apparent that a few pathways, such as those involved in cell wall biosynthesis pathways and lactose and galactose degradation I (LACTOSECAT-PWY) were enriched in *Tetragenococcus*-dominated microbiome (Figure 4B). However, a diverse array of predicted pathways were enriched in non-*Tetragenococcus*-dominated microbiome, especially those involved in purine biosynthesis, amino acid metabolism and vitamin/cofactor/carrier biosynthesis. Purine and amino acid biosynthesis are implicated with organoleptic quality of fermented foods. Therefore, we attempted to look within the non-*Tetragenococcus*-dominated microbiome for predicted contributions of individual genus to these two pathways. We did not observe changes in predicted enriched pathways across sampling batches.

**Figure 4:**
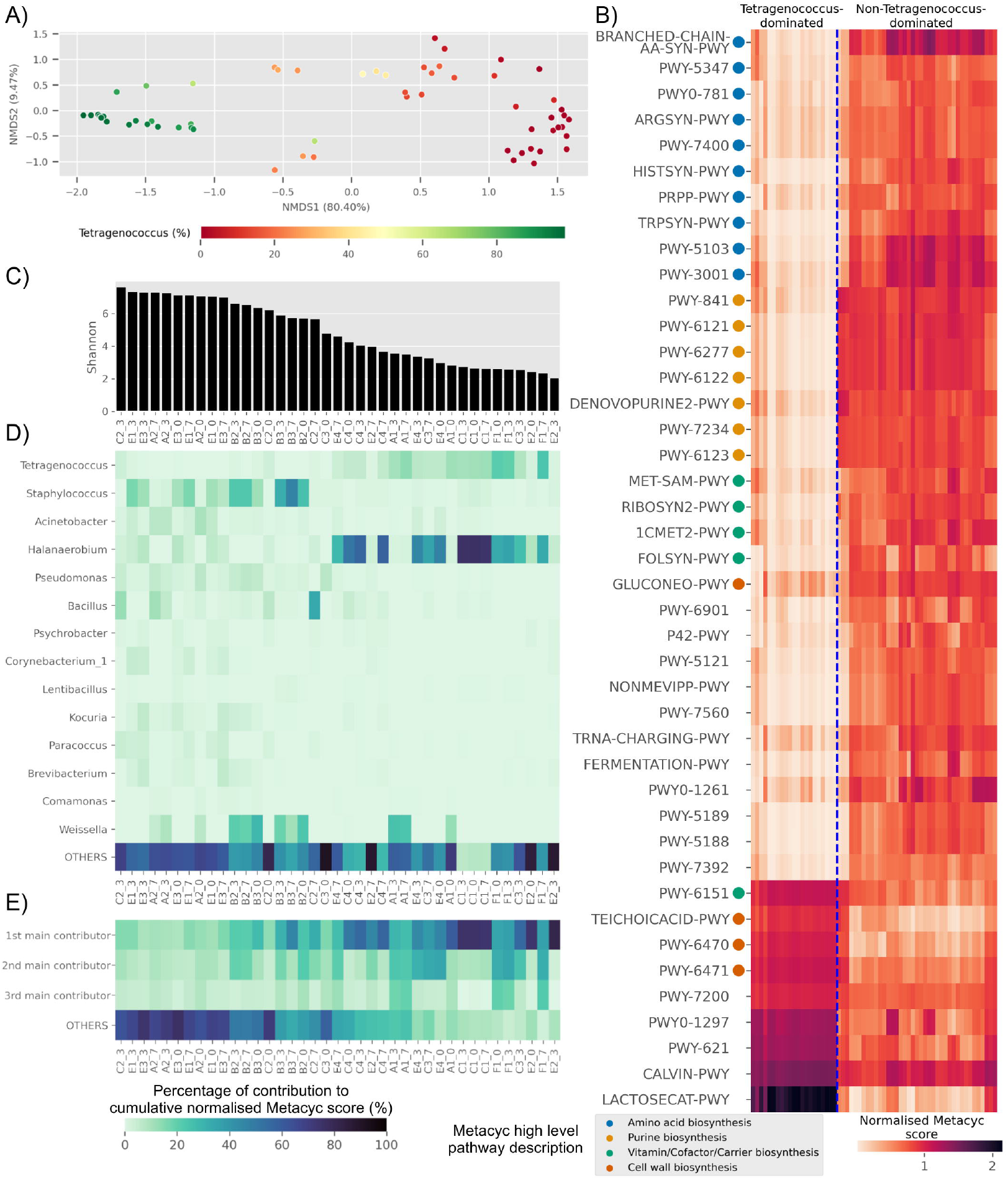
A) Multidimensional scaling plot based on overall predicted pathways for each samples. Sample are coloured based on relative abundance of *Tetragenococcus*, B) Heatmap of differential pathways between *Tetragenococcus*-dominated and non-*Tetragenococcus*-dominated samples based on normalised METACYC score. Pathways labels are based on shortened pathway labels from the Metacyc database, C) Shannon index, sorted by value. Only non-*Tetragenococcus*-dominated samples are shown, D) Heatmap of pooled contribution towards predicted organoleptic pathway per genus of prevalence >= 50% per sample. Only non-*Tetragenococcus*-dominated samples are shown, E) Pooled contribution towards predicted organoleptic pathway of top three most abundant genus per sample. Only non-*Tetragenococcus*-dominated samples are shown.

Aside from *Tetragenococcus, Staphylococcus, Halanaerobium, Bacillus* and *Weissella*, other prevalent genera did not appear to contribute much towards purine and amino acid biosynthesis, relative to the cumulative contribution of the remaining lesser genera (OTHERS) (Figure 4D). By focusing on the top three most abundant genera per sample regardless of taxonomic prevalence, purine and amino acid biosynthesis in samples of low Shannon index (Figure 4C) were contributed by the dominant genus (Figure 4E). This includes samples without high prevalence genus such as samples from bottle E2 (Figure 4D). This was not observed in samples with high Shannon diversity index. For instance, samples from bottles E1 and E3 which have relatively high Shannon scores possess high percentage of contribution from the remaining genera instead of the top three most abundant genera per sample (Figure 4E).

There are several predicted pathways which are enriched in non-*Tetragenococcus*-dominated samples, such as L-methionine biosynthesis (transsulfuration) (PWY-5347), L-histidine biosynthesis (HISTSYN-PWY), flavin biosynthesis I (bacteria and plants) (RIBOSYN2-PWY) and superpathway of tetrahydrofolate biosynthesis and salvage (FOLSYN-PWY), which nutrients are only to be obtained through diet.

### 3.4 Microbial abundance interaction network

After prevalence filtering of the ASV to be present in at least 50% of samples, there were a total of 71 nodes in the predicted microbial interaction network (Figure 5A), 73.7% were positive associations, and 26.3% were negative associations. Several isolated sub-clusters formed such as a small cluster consisting of four nodes (ASV4, ASV7, ASV10 and ASV11); all four nodes were assigned as *Tetragenococcus halophilus* subsp. Halophilus. The 16S copy number predicted from these ASVs are just one copy per ASV, which implies that the ASVs possibly belong to closely related *T. halophilus* strains. The isolation of this cluster might imply a lack of interaction between *Tetragenococcus halophilus* subsp. Halophilus with the rest of the Budu inhabitants. The nodes are not fully connected with each other, possibly because of the conditional independence implemented in SPIEC-EASI to prevent spurious links. Positive associations were predicted for edges ASV4-ASV7 and ASV7-ASV11. However, a negative association formed between ASV4 and ASV10.

**Figure 5:**
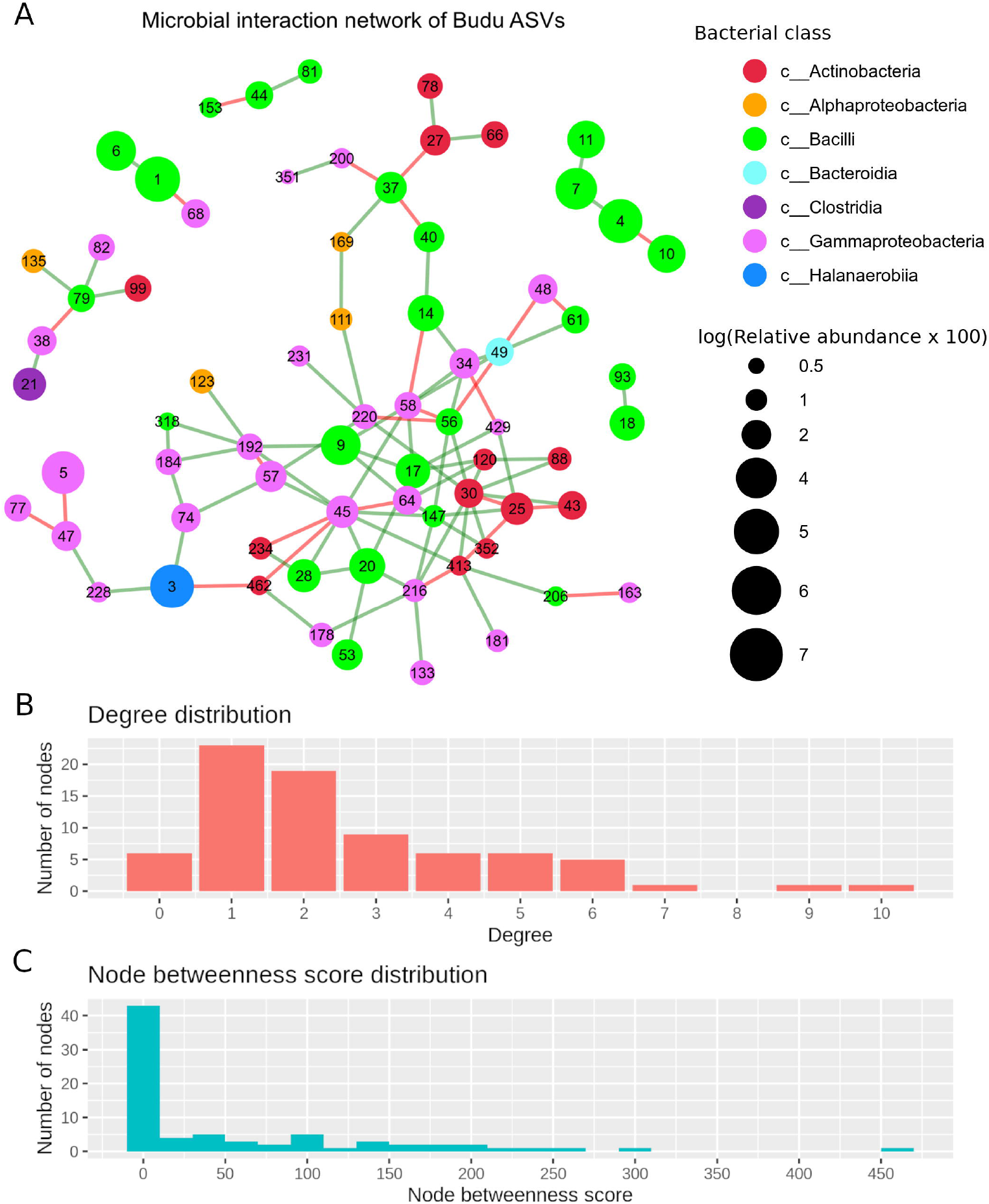
A) Predicted interaction network based on Meinshausen-Bühlmann neighborhood selection model. Nodes represent ASVs, coloured based on class taxonomic assignment, and node size was given based on relative abundance, scaled logarithmically. Green edges indicate positive interaction, while red edges indicate negative. To shorten the node label, the letters “ASV” was removed from the node label, B) Distribution of degree distribution, C) Distribution of degree betweenness scores

Several nodes of similar taxonomic assignment were predicted to interact with each other such as that of the *T. halophilus* subsp. Halophilus sub-cluster as well as ASV1-ASV6 (*Tetragenococcus muriaticus*), ASV47-ASV77-ASV228 (*Acinetobacter*), ASV20-ASV28-ASV53 (*Staphylococcus*), ASV216-ASV133 (*Psychrobacter*) and ASV27-ASV66 (*Kocuria*). Co-abundance between these nodes were expected due to 16S copy numbers of slight nucleotide variations originating from the same bacterial genome. There are also patterns of network assortativity, which was most apparent for the *Actinobacteria* nodes surrounding ASV30 which is assigned as *Brevibacterium*. ASV30 formed a number of positive links with *Corynebacterium* nodes which belonged to the Actinobacteria class.

Degree centrality and betweenness centrality were two node centrality measures computed from the predicted interaction network. Majority of the nodes had very low degrees and possessed low betweenness centrality scores. ASV45 (Enterobacteriaceae) boasted the highest degree centrality (degrees = 10) and betweenness centrality (score = 466.72). *Tetragenococcus* nodes have very low node degrees and betweenness. Only one edge that formed between *Tetragenococcus* nodes and non-*Tetragenococcus* nodes (ASV1 *T. muriaticus* with ASV68 *Pseudomonas*).

## 4 DISCUSSION

Fermented fish sauce is a very popular food condiment in the South East Asian region. The fermentation from raw fish is largely driven by its microbiome because the high salt concentration in fish sauce inhibits the activity of endogenous fish enzymes like cathepsins, but activates halophilic bacterial proteinases (Tungkawachara et al., 2003)□. Since then, several microbiome-based research were done on fermented fish sauce in Korea, which has revealed insights into the role of the microbiome in industrial fish sauce fermentation as well as fish sauce quality (Jung et al., 2016; Lee et al., 2014). However, current studies on Budu so far are based on culture-based methods which have limited taxonomic breadth; it has not been studied yet to a degree that would enable understanding of the overall microbiome and its potential function. This is needed because its consumption can present as a potential health risk due to its high purine and histamine content (Mohd et al., 2011; Rosma et al., 2009). Due to the rich purine content in anchovies, (2,258.91 mg/kg) (Li et al., 2019), there is a concern that consumption of Budu could present a risk to higher gout incidence in Malaysia (Paul and James, 2017). Also, based on Rosma et al. (2009), seven of twelve Budu samples from twelve different producers contained histamine content exceeding 50 mg/100 g sample, which is the FDA limit for histamine consumption (FDA, 2011)□. Unchecked consumption of histamine-rich food can lead to scombroid food poisoning, the symptoms which are breathing difficulties and irregular heartbeats (Tortorella et al., 2014; Wilson et al., 2012). By determining the microbial diversity and composition of Budu, we hope to elucidate the relationship between the microbiome in Budu and and Budu quality, which can later translate to enhancement of organoleptic and nutritional value of Budu.

Clear inconsistencies of Budu microbiome were apparent within the same brand in terms of phyla composition, reiterated by the huge distances in PCoA as well as differences in alpha diversity indices. This suggests that most Budu were produced under uncontrolled conditions between production batches which may lead inconsistencies in the microbiome (Jung et al., 2016). This is also a major obstacle to commercial production with uniform, premium quality. From a food production point of view, inconsistent microbial diversity might interpret to different fermentation performances and quality variations. However, it is amusing to see a wider taxonomic diversity of bacteria that can thrive in Budu and potentially contribute to its quality. Future research should include the microbiota composition to predict organoleptic quality of the fish sauce. Our results also highlight the utility of sequencing as a potential tool to assess consistency in fermentation food production.

*Tetragenococcus* is a tetrad-forming coccus, gram positive halophilic lactic acid bacteria. It was previously isolated from high-salt fermented foods such as soy sauce, squid liver sauce (Satomi et al., 1997), dried fish, salted seafood (Kim and Park, 2014), fish sauce (Thongsanit et al., 2002) and fermented fish paste (Marui et al., 2015). Despite being the genus with the highest average relative abundance and prevalence, previous studies did not report of its isolation from Budu. Unsuccessful isolation of *Tetragenococcus* from Budu might be attributed to limited sampling or culturing conditions unsuitable to *Tetragenococcus*, showing the caveats of culturing-based methods. Despite its prevalence in numerous fermented food, to the best of our knowledge, *Tetragenococcus* was never found in the natural environment, neither found in fish gut nor saltern sources (Egerton et al., 2018). *Tetragenococcus* was highly abundant in most Budu samples but was very low or absent in other samples. We also identified two *Tetragenococcus* species which were *T. halophilus* (6 ASVs) and *T. muriaticus* (2 ASVs) (Kobayashi et al., 2000). Based on 16S copy prediction, *Tetragenococcus* nodes possess one copy of 16S sequencing. Therefore, the different ASVs found in our samples suggest the presence of different strains of *Tetragenococcus*, including *T. halophilus* and *T. muriaticus*. The two species were reported to grow optimally at different pH levels and salt concentrations and utilize different types of sugar (He et al., 2016; Kobayashi et al., 2004, 2000; Udomsil et al., 2010). *Tetragenococcus muriaticus* is known to release histamine and cadaverine, while *Tetragenococcus halophilus* is known for the ability to reduce histamine and cadaverine formation (Kim et al., 2019; Zaman et al., 2010). *T. halophilus* was also reported to contribute towards probiotic properties (Kuda et al., 2014). It is not yet understood how *Tetragenococcus* dominated a microbiome, or if it is possible to control its abundance – the consequences would include manipulation of the microbiome to enrich for desirable metabolites or microbial processes.

The Budu microbiome composition was generally consistent post-fermentation. However, it was not stagnant, as shown by the significant increase of *Chromobacterium* in the last sampling batch in brand E, followed by brand D. Since *Chromobacterium* was not detected in first and second batches, it is also possible that the genus was introduced from the surrounding into the samples while sampling from the previous sampling batches. Brand D and brand E did not exclusively share common ingredients. The two brands are also different in terms of microbiome composition, therefore we are unsure of what factors caused the emergence of *Chromobacterium* in the two brands. *Chromobacterium* was previously found in environmental samples (Kämpfer et al., 2009; Menezes et al., 2015) as well as food samples such as vegetables, cheese and seafood samples (Koburger and May, 1982). Bacteremia-causing *Chromobacterium* species were also reported (Kaufman et al., 1986; Parajuli et al., 2016). Due to the lack of genomic sequences and analyses of the *Chromobacterium* species (Santos et al., 2018), future research should address this gap of knowledge and identify the ecological significance of this genus in fermented food.

The microbiome determines the metabolism taking place in Budu, which was clearly predicted between *Tetragenococcus*-dominated and non-*Tetragenococcus*-dominated samples. The latter was predicted to be enriched with biosynthesis of amino acids and purine, which were known to contribute towards the quality of fermented foods in terms of pleasant organoleptic characteristics. For instance, umami flavour in food is given by macromolecules which are amino acids such as glutamate and aspartate as well as purine such as inosinic acid and guanylic acid (Zhao et al., 2019)□. These pathways are also implicated with health conditions. For instance, metabolites enriched in Budu such as histamine and purine compounds are implicated with health conditions such as scromboid poisoning and gout respectively (Paul and James, 2017; Tortorella et al., 2014)□. The predicted functions of the microbiome demonstrates the functional potential of the microbial community but does not necessarily translate to the activities taking place in the community. To do so, more omics data are required from genomics (transcriptomics) and also associated metabolomics data.

Unfortunately, these information were not yet within the scope of this current study. Thus, future studies can focus more on the influence of amino acid and purine biosynthesis on Budu quality while accounting for the microbial diversity as seen in our samples. A recent study demonstrated that amino acids and biogenic amines increased in fermented fish sauce after a period of a year post-fermentation and was also influenced by storage conditions such as temperature (Joung and Min, 2018)□. Another study showed that prolonged fermentation of the fish can also lead to conversion of glutamic acid available in the fish protein into more purine (Tungkawachara et al., 2003). However, we did not observe changes in predicted enriched pathways across sampling batches.

Microbes exist in communities and interact with each other through mechanisms such as production of lactic acid, the release of antimicrobials, quorum sensing, and cross-feeding. The ability to broadly predict such relationships using relative abundance information from 16S sequencing is alluring, thus attempts on modeling microbial interactions were done in the form of microbial interaction network (Deng et al., 2012; Faust et al., 2012; Friedman and Alm, 2012; Kurtz et al., 2015). From our constructed interaction network, we observed small isolated clusters, including a four-nodes cluster comprising ASV4, ASV7, ASV10 and ASV11, and all the nodes were assigned to *Tetragenococcus halophilus* subsp. Halophilus. No interactions were predicted between *T. halophilus* nodes with other taxa. Known interaction of *Tetragenococcus halophilus* was reported in soy sauce fermentation in the form of antagonism with *Zygosaccharomyces rouxii* which is a yeast (Devanthi et al., 2018). *T. muriaticus* was another *Tetragenococcus* species that was detected in our study; the ASVs assigned to *T. muriaticus* are ASV1 and ASV6 and their nodes are linked in the network. There was only one non-*Tetragenococcus* node that linked with *Tetragenococcus* node, which was ASV68, assigned as *Pseudomonas*. Since the network is based on relative abundance information, the lack of connections that formed between *Tetragenococcus* nodes and non-*Tetragenococcus* nodes could mean either the interaction network prediction could not model in situations where ASVs that are highly dominant or that *Tetragenococcus* form only a few interactions with other taxa. Known interaction of *Tetragenococcus halophilus* was reported in soy sauce fermentation in the form of antagonism with *Zygosaccharomyces rouxii* which is a yeast (Devanthi et al., 2018). The interpretation of microbial interaction network and its application in a biological context is still limited. For example, Bacillus is both abundant and prevalent in our study. However, the high salinity of Budu (>15%) (Mohamed et al., 2012) was known to inhibit growth of *B. subtilis* of marine origin (Katja et al., 2014)□. Thus, it could be that the 16S sequence of *Bacillus* was amplified from spore particles (Fukui et al., 2012). Research in microbial interaction network is also hampered by the lack of gold standard, thus it is difficult to disentangle true interactions from strong environmental effect. Budu easily lends itself as a tractable microbial system which is replicable and manipulable, and it is cheap, making it a potential gold standard for this benchmark current and future microbial interaction network models (Wolfe, 2018; Wolfe and Dutton, 2015).

## 5 CONCLUSIONS

The microbiome of some brands of Budu are inconsistent, which possibly suggests Budu fermentation was done under uncontrolled conditions between production batches. With the increasing popularity of fermented food and the descending sequencing cost, microbiome assessment through DNA sequencing is now a viable and cost effective method in quality assurance of fermented food. We also discovered abundant and prevalent genera of Budu, some of which were not previously found in previous studies on Budu which were based on culturing methods. Albeit not reported in prior culture-dependent studies on Budu, *Tetragenococcus* was the most abundant genus in our sample collection, and the two of its species that were detected were *Tetragenococcus muriaticus* and *Tetragenococcus halophilus*. While *Tetragenococcus* was a common dweller in fermented fish sauce and other fermented seafood, some of our Budu samples were devoid of the genus. This points out how Tetragenococcus is well adapted to fermentation environment but also points towards a greater diversity of bacteria that are as capable as *Tetragenococcus* in fulfilling its role in fermentation. Non-*Tetragenococcus* dominated samples were predicted to be enriched with metabolic pathways associated with amino acid biosynthesis and purine biosynthesis, macromolecules that were attributed to organoleptic properties as well as nutrition. This opens up towards exploration of a wider array of microbes as a candidate starter culture of Budu to improve Budu quality and safety. We also detected *Chromobacterium* as the only genera that was significantly increased in the last sampling batch, though its ecological role and its potential source was not yet clear. We also attempted to model the microbial interaction using 16S abundance data, although its interpretation is still quite limited.

## 6 DATA ACCESS

All FastQ raw data can be accessed through SRA accession SRP193353, or through NCBI BioProject under BioProject ID: PRJNA534025. Supplementary files are accessible through the Mendeley Dataset http://dx.doi.org/10.17632/6jhvcgh6fm.1

## 9 ACKNOWLEDGMENTS

We are very grateful towards Wan Nur Afiq and Wan Nur Fariees Fitrie for providing Budu samples from Kelantan. We want to thank Monash University Malaysia Genomics Facility for financially supporting the project and providing computational resources. We are also thankful to Amazon Web Services ASEAN Public Sector, particularly Sammy Lock and Shazli Mohd Ghazali for provision of proof of concept (POC) credits which we used for computationally intensive microbial interaction prediction. Finally, we express our gratitude to Shu Yong Lim and Sze Mei Lee for reviewing the manuscript and suggest changes.

## 10 CRediT AUTHOR STATEMENT

Muhammad Zarul Hanifah: Data curation; Formal analysis; Investigation; Methodology; Resources; Visualization; Roles/Writing -original draft

Han Ming Gan: Conceptualization; Funding acquisition; Methodology; Project administration; Resources; Supervision; Writing -review & editing

## 11 Figure AND TABLE CAPTIONS

** All Figures are to be treated as coloured.

**Supplementary Figure 1:**
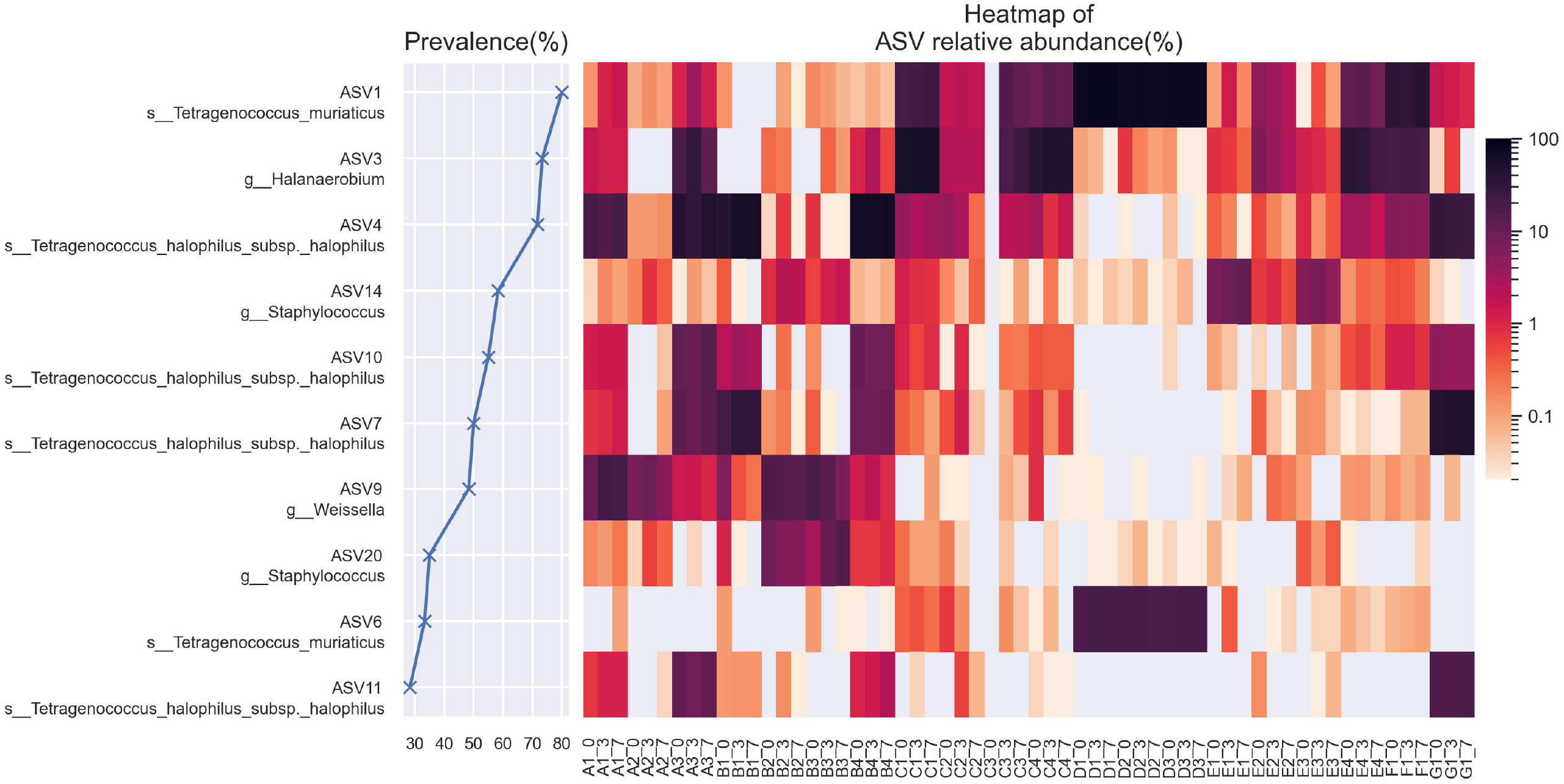
Prevalence and abundance per sample for dominant ASVs (ASVs displaying maximum abundance of >10% in at least one sample)

**Supplementary Figure 2:**
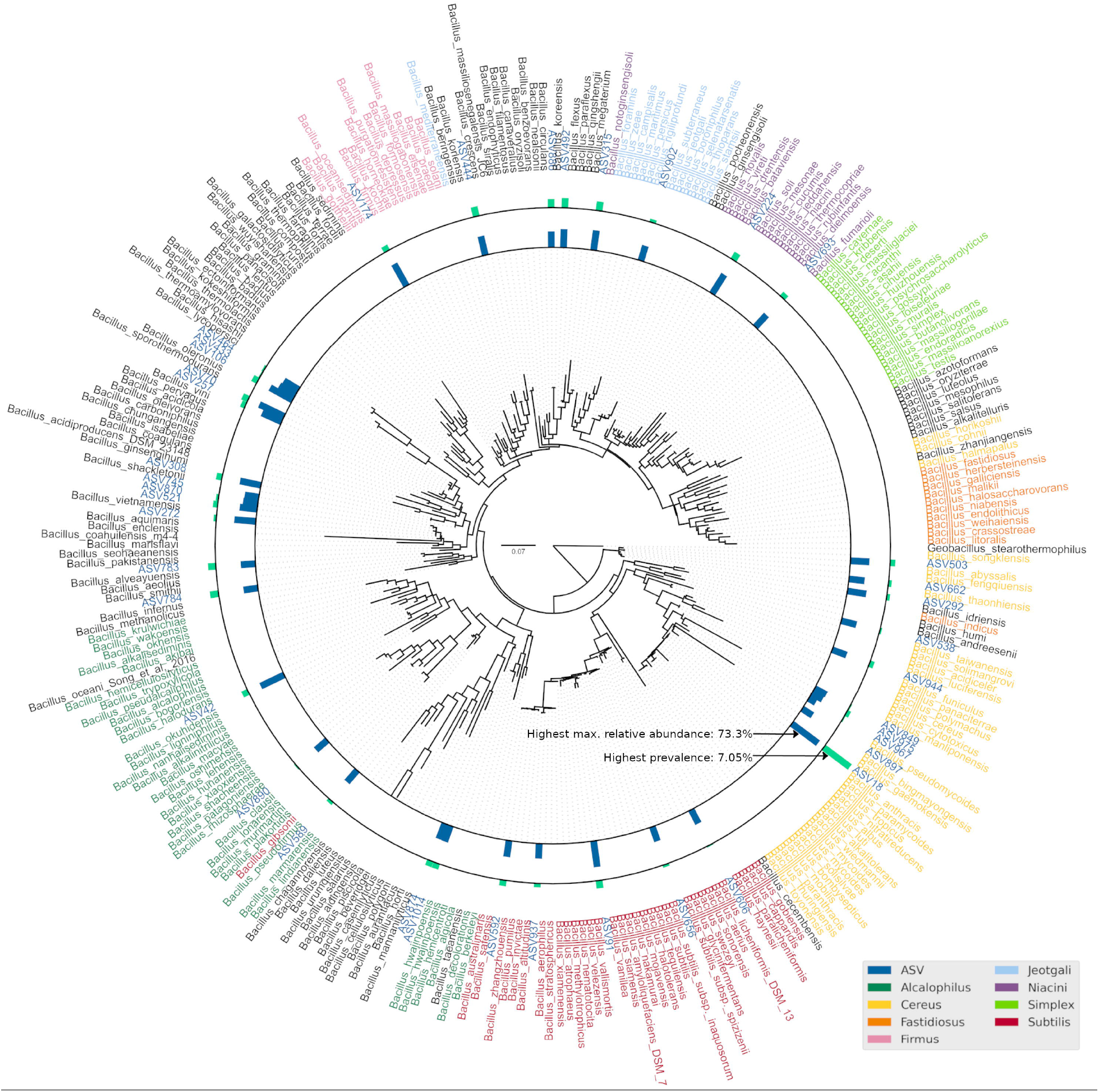
Maximum likelihood phylogenetic tree of representative *Bacillus* sequences from SILVA and RDP databases together with ASVs assigned as *Bacillus*. Bar plots represent prevalence (blue, max. = 73.33 %) and maximum relative abundance of ASV (green, max. = 7.05%). ASV18 was the most abundant and prevalent *Bacillus* ASV. Leaf nodes are colored according to Patel and Gupta (Patel and Gupta, 2020)□ *Bacillus* clade designation. The phylogenetic tree was rooted to *Geobacillus stereothermophilus* as outgroup.

**Supplementary Figure 3:**
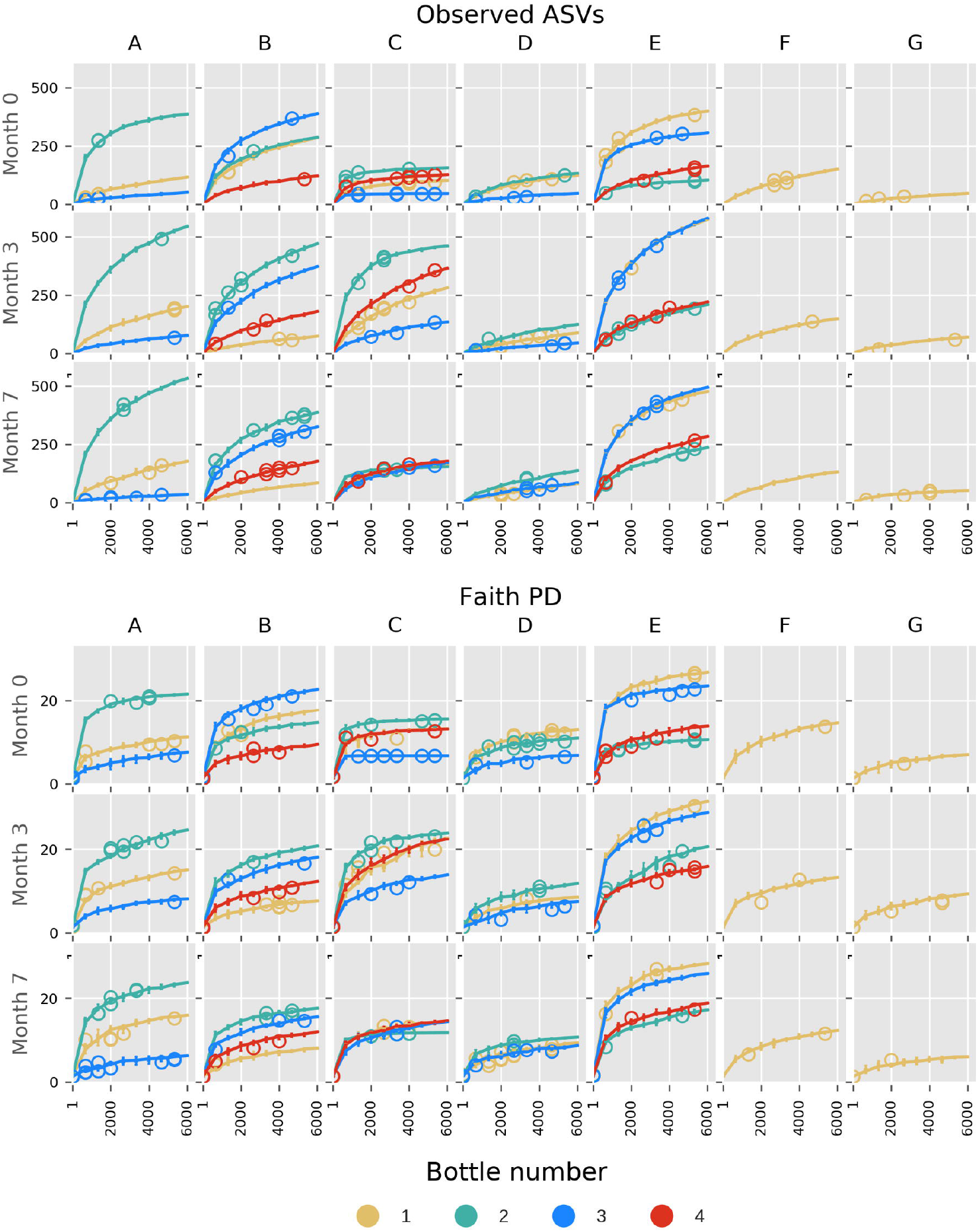
Alpha rarefaction curves based on observed ASVs and Faith Phylogenetic Diversity, faceted by brand and sampling period. The interquartile ranges are shown. Circles indicate outlier alpha diversity calculations

**Supplementary Figure 4:**
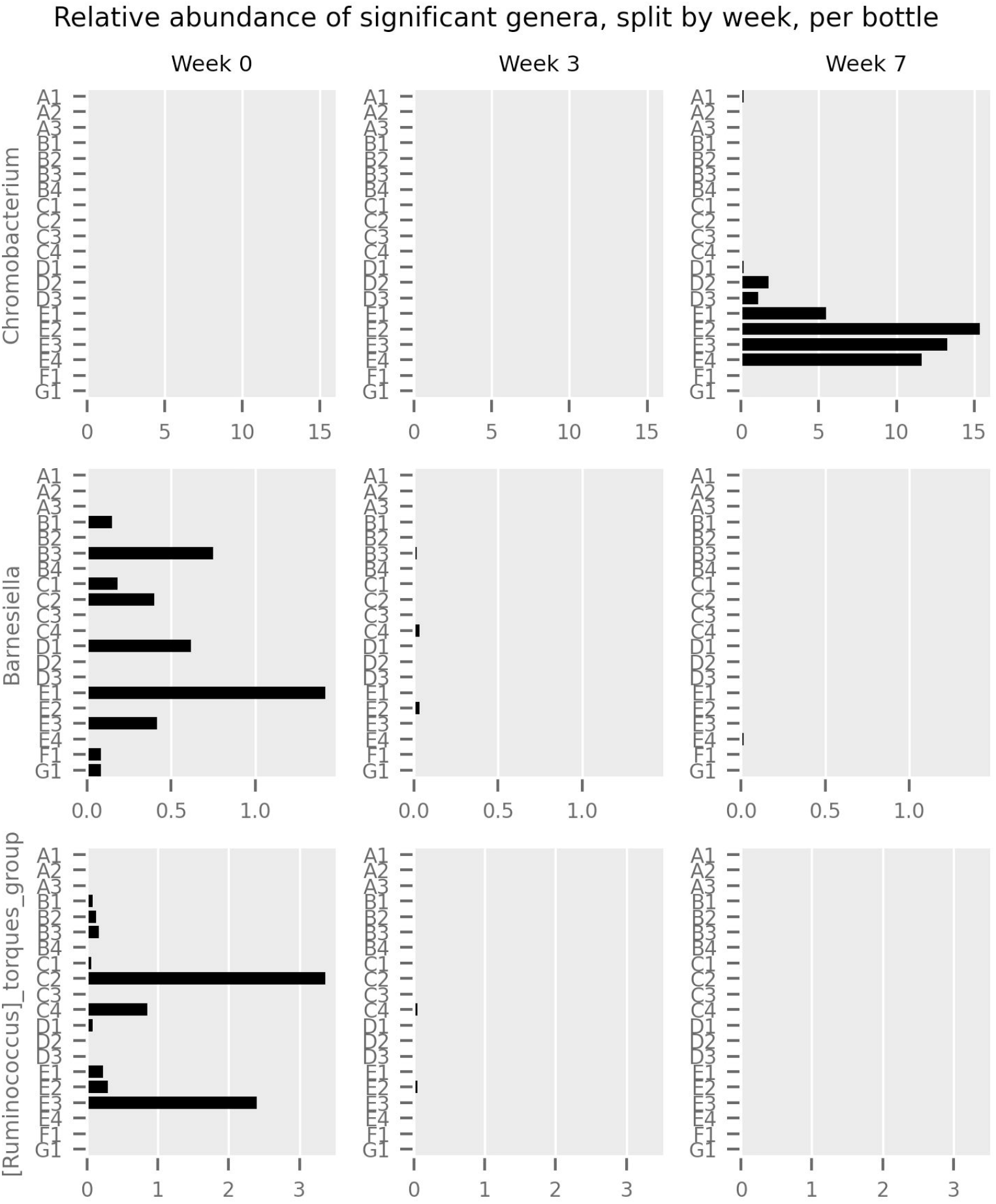
*Relative abundance of Chromobacterium, Barnesiella* and *[Ruminococcus] torques* group across sampling periods per sample

## 12 SUPPLEMENTARY MATERIAL CAPTIONS

Supplementary File 1: ASV sequences in fasta format

Supplementary File 2: Taxonomic assignment table of individual ASVs using q2-classifier

Supplementary File 3: Observation table of all samples after filtering out the ASVs of non-microbial, rarefied at 6000 reads per sample

Supplementary File 4: Predicted functional table based on EC. The table is stratified by ASV Supplementary File 5: Predicted pathway table based on Metacyc database. The table is stratified by ASV

